# ACDC: Analysis of Congruent Diversification Classes

**DOI:** 10.1101/2022.01.12.476142

**Authors:** Sebastian Höhna, Bjørn T. Kopperud, Andrew F. Magee

**Affiliations:** GeoBio-Center LMU, Ludwig-Maximilians-Universität München, Richard-Wagner Straße 10, 80333 Munich, Germany; Department of Earth and Environmental Sciences, Paleontology & Geobiology, Ludwig-Maximilians-Universität München, Richard-Wagner Straße 10, 80333 Munich, Germany; Department of Human Genetics, University of California, Los Angeles, 90095, U.S.A.

**Keywords:** Birth-death models, macroevolution, diversification rates, identifiability, congruence class

## Abstract

1. Diversification rates inferred from phylogenies are not identifiable. There are infinitely many combinations of speciation and extinction rate functions that have the exact same likelihood score for a given phylogeny, building a congruence class. The specific shape and characteristics of such congruence classes have not yet been studied. Whether speciation and extinction rate functions within a congruence class share common features is also not known.

2. Instead of striving to make the diversification rates identifiable, we can embrace their inherent non-identifiable nature. We use two different approaches to explore a congruence class: (i) testing of specific alternative hypotheses, and (ii) randomly sampling alternative rate function within the congruence class.

3. Our methods are implemented in the open-source `R` package `ACDC` (https://github.com/afmagee/ACDC). `ACDC` provides a flexible approach to explore the congruence class and provides summaries of rate functions within a congruence class. The summaries can highlight common trends, i.e. increasing, flat or decreasing rates.

Although there are infinitely many equally likely diversification rate functions, these can share common features. `ACDC` can be used to assess if diversification rate patterns are robust despite non-identifiability. In our example, we clearly identify three phases of diversification rate changes that are common among all models in the congruence class. Thus, congruence classes are not necessarily a problem for studying historical patterns of biodiversity from phylogenies.

## 1 Introduction

In macroevolution, one prominent avenue of research is to estimate macroevolutionary rates of diversification from molecular phylogenies (Ricklefs 2007; Morlon 2014). Specifically, many studies are interested in inferring time-varying diversification rates. Time-varying diversification rates are used to study monotonous slowdowns/increases using continuous functions (*e.g.*, Rabosky 2006; Morlon *et al.* 2011; Höhna 2014), abrupt shifts in diversification rates (*e.g.*, Stadler 2011; May *et al.* 2016; Magee *et al.* 2020), mass extinction (*e.g.*, Höhna 2015; May *et al.* 2016; Culshaw *et al.* 2019; Magee & Höhna 2021) and correlations to environmental factors (*e.g.*, Condamine *et al.* 2013; 2019; Palazzesi *et al.* 2022). Unfortunately, time-varying diversification rates are not identifiable when estimated from time-calibrated phylogenies, while allowing for any continuous diversification rate function (Kubo & Iwasa 1995; Louca & Pennell 2020). That is, infinitely many combinations of speciation and extinction rate functions, summarized within a *congruence class*, result in the same likelihood given a phylogenetic tree (Kubo & Iwasa 1995; Louca & Pennell 2020).

However, the existence of infinitely many equivalently likely rate functions does not imply that one cannot draw any general conclusions. We do not know yet which diversification rate functions are within a congruence class, and if these diversification rate function share some specific features (e.g., rate changes at the same time). If we obtained estimates of diversification rates for our study group, then we could be interested in exploring all or a sample of diversification rates included in the congruence class to identify shared features. Furthermore, we could test if different specific diversification rate scenarios are included within a congruence class, for example, if a model with exponentially increasing/decreasing diversification rates is included in the congruence class. Similarly, we could test if models with rate shifts at specific times, corresponding to alternative hypotheses, are included in the congruence class.

Here we provide the `R` package `ACDC` (Analysis of Congruent Diversification Rates) that (1) converts between models within the same congruence class, (2) explores the full congruence class, and (3) shows common trends among models within the same congruence class. Conversion between congruent models is useful if a researcher wants to explore alternative hypotheses, for example, “what if the extinction rate was not constant but instead exponentially increased through time?” Full exploration of the congruence class can highlight general features of a congruence class, for example, if a researcher has estimated a given pattern of diversification rates and wants to know if all models within the congruence class show a certain trend (e.g., a rate shift at time *t*). Finally, all explored models can be analyzed to show common patterns of diversification rate increases and decreases. With our `R` package `ACDC`, researchers can test which patterns are robust to the congruence class.

## 2 Theory and Usage

In this section we provide the theory and our approach as well as how to use `ACDC`. We start by explaining how `ACDC` processes any diversification rate function to construct the congruence class. Next, we explain how `ACDC` obtains alternative (congruent) matching pairs of speciation and extinction rates functions. Then, we show how specific hypotheses can be tested and how to explore the congruence class using diversification rate functions generated from a process or a distribution. Finally, we demonstrate how alternative diversification rate models within the congruence class can be summarized to assess for shared trends.

### 2.1 Linear interpolation

Congruence classes are derived for arbitrary continuous functions of speciation and extinction rates (Louca & Pennell 2020). Unfortunately, it is not feasible to always work with arbitrary continuous functions because the derivative is unknown. Instead, we use a piecewise linear approximation of arbitrary user-defined rate functions to enable the exploration of congruence classes (Fig. 1). This approximation should be unproblematic for representing existing models, especially since piecewise constant diversification models are often used for diversification rate inference (Stadler 2011; May *et al.* 2016; Magee *et al.* 2020; Magee & Höhna 2021). Moreover, the piecewise linear approximation can become arbitrarily close to any continuous function if the number of pieces is sufficiently large. We show the effect of the number of pieces used on the accuracy of the approximation in the Supplementary Material.

**Figure 1:**
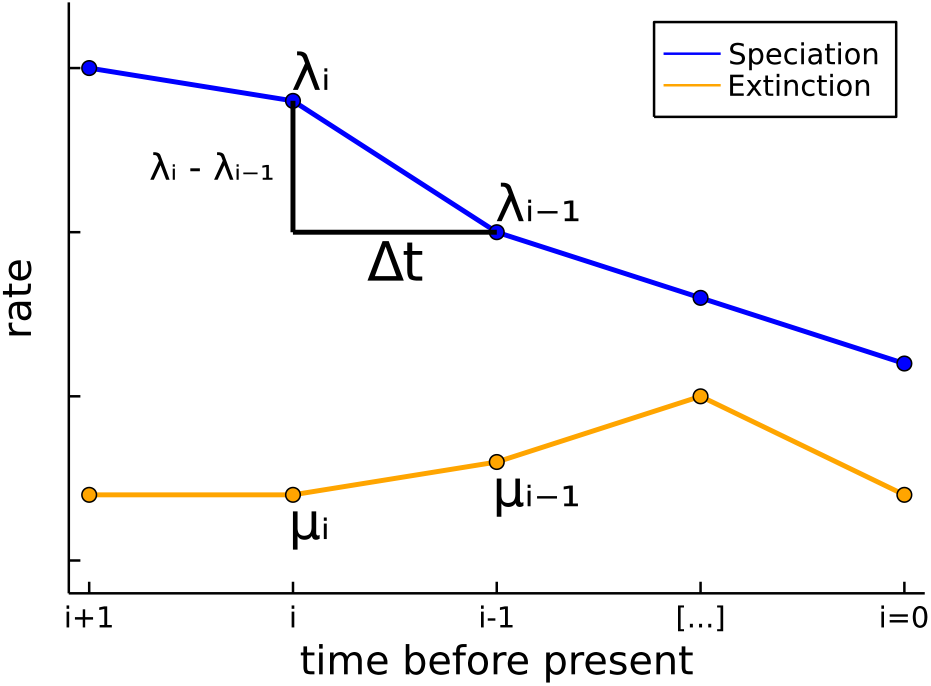
Schematic of the piecewise linear rate functions. The piecewise linear function shows how any continuous function can be approximated if sufficiently many linear components are used. We use the index *i* = 0 for the present, and increasing toward the past. The finite difference *λ_i_ − λ*_*i*−1_ per time interval Δ*t* represents the slope of the rate function. The finite difference is important to analytically compute the slope (i.e., the derivative) of the rate function at any time.

We assume a grid of *n* + 1 times, *t*_0_*, t*_1_*, …, t_n_*, with constant spacing Δ*t*. This grid typically spans between 0 and the root age of the phylogenetic tree. The piecewise linear speciation rate approximation is defined by linear interpolation of the vector ****λ**** = *λ*_0_*, …, λ_n_*. Specifically, as depicted in Fig. 1, the (interpolated) speciation rate within the interval *t_i_*_−1_ < *t* ≤ *t_i_* is

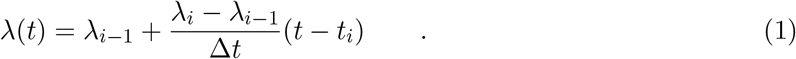

The extinction rate approximation is defined analogously.

As an empirical example, we estimated speciation and extinction rates for the primates phylogeny from Springer *et al.* (2012) using a horseshoe Markov random field (HSMRF) prior distribution (Carvalho *et al.* 2010; Magee *et al.* 2020) as implemented in `RevBayes` (*Höhna et al.* 2016). The specific details about the data set and Markov chain Monte Carlo settings are not important for this study, but can be found at https://revbayes.github.io/tutorials/divrate/ebd.html. We include the samples from the posterior distribution in our package for convenience. We will use this example to showcase how to explore the congruence class with `ACDC`. In `R`, we can set up the piecewise linear rate functions as follows.

~~~
library(ACDC)
data(primates_ebd)
lambda <- approxfun(primates_ebd$time, primates_ebd$lambda)
mu <- approxfun(primates_ebd$time, primates_ebd$mu)
~~~

### 2.2 Constructing the Congruence Class

The central idea in `ACDC` is to construct the congruence class given a speciation and extinction rate function. A congruence class is fully specified by either the pulled net-diversification rate *r_p_*(*t*) and the speciation rate at the present (*λ*_0_), or the pulled speciation rate *λ_p_*(*t*). That is, any combination of speciation and extinction rate function that result in the same pulled net-diversification rate *r_p_*(*t*) and speciation rate at the present (*λ*_0_), or pulled speciation rate *λ_p_*(*t*), belong to the same congruence class (Louca & Pennell 2020). We can therefore set up the congruence class by constructing the pulled net-diversification rate *r_p_*(*t*). The pulled net-diversification rate is defined as (Louca & Pennell 2020)

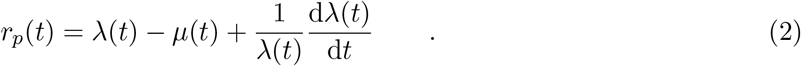

Since we are using piecewise linear speciation and extinction rates (Fig. 1), we obtain (at the interval times) the analytical solution for the pulled net-diversification rate as

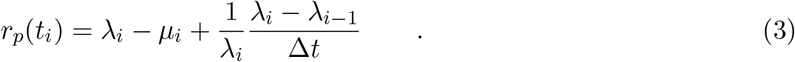

The equation for the pulled speciation is (Louca & Pennell 2020):

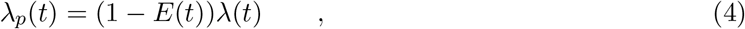

where *E*(*t*) is the probability that the lineage observed at time *t* goes extinct before the present. Currently we only use the pulled speciation rate for plotting purposes.

We use the pulled net-diversification rate to construct the congruence class because it simplifies the equations. Continuing with the primates data example, we can construct the congruence class in `ACDC` as follows.

~~~
times <- seq(0, max(primates_ebd$time, length.out = 1000)
my_model <- create.model(lambda, mu, times)
~~~

We use a fine grid of one thousand time points to improve the precision of the calculations. Then, we can plot the diversification rates together with their pulled counterparts in `ACDC` using `plot(my_model)` (Fig. 2). Studying the pulled speciation and pulled net-diversification rates itself can highlight aspects of the congruence class (Helmstetter *et al.* 2021).

**Figure 2:**
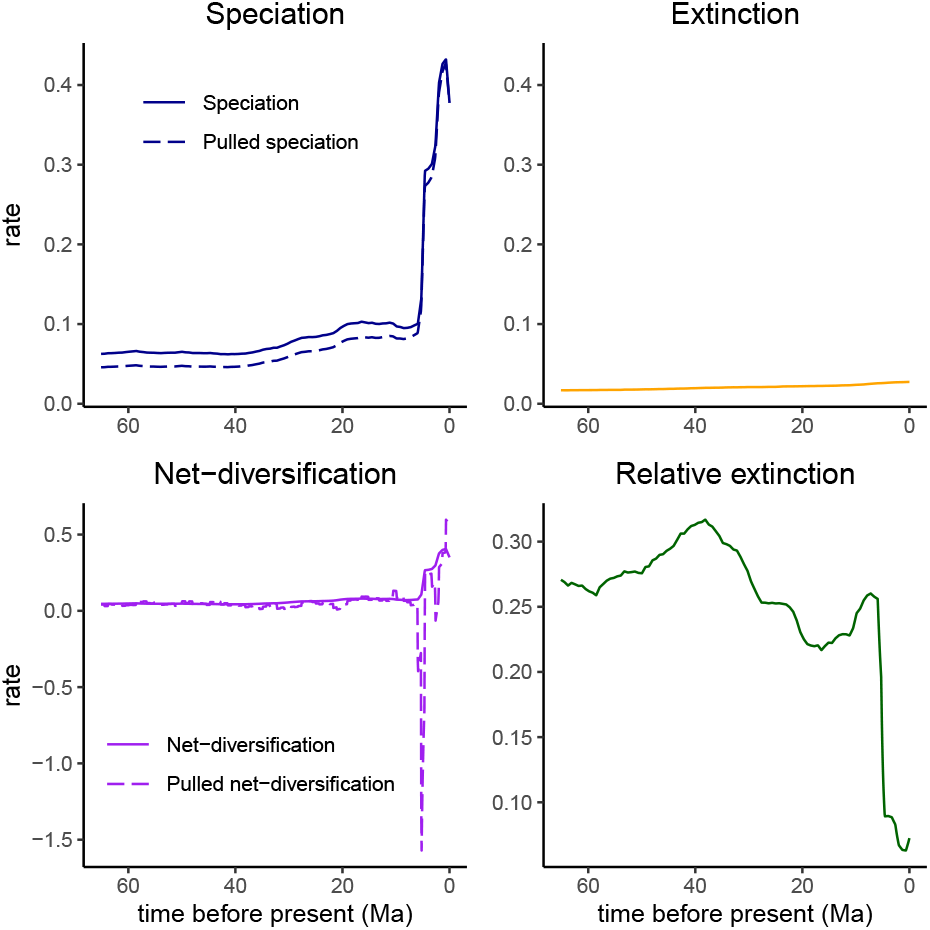
Estimated diversification rates for the primates phylogeny by Springer *et al.* (2012) using an episodic birth-death model with a horseshoe Markov random field prior (Magee *et al.* 2020) in `RevBayes` (Höhna *et al.* 2016). The original rates define the *reference* model in `ACDC` (solid lines). `ACDC` automatically computes the pulled speciation rate as well as the pulled net-diversification rate (dashed lines), which characterize the congruence class.

### 2.3 Transforming speciation and extinction rates

A researcher can either provide another speciation rate function (or extinction rate function) and `ACDC` computes the corresponding extinction rate function (or speciation rate function, respectively) so that the new pair also belongs to the same congruence class. If a user provides a new extinction rate function, and wishes to know the speciation rate, we can compute it as:

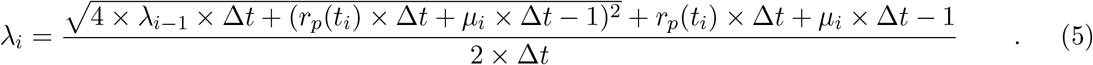

We show in the Supplementary Material how Eq. (5) is derived from Eq. (3). We note that *λ*_0_ is equal for all models in the congruence class. Conversely, if a user provides a new speciation rate function, then we can solve Eq. (3) for *μ* and get

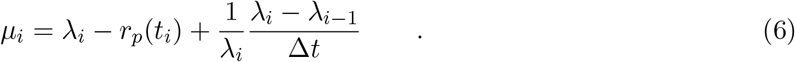

These two transformations allow us to explore the congruence class. We only need to propose an alternative speciation rate function, or an alternative extinction rate function, and then compute their counterpart for the new model to be within the congruence class.

### 2.4 Exploring congruent models for specific hypotheses

A first option to explore the congruence class is to test for specific alternative hypotheses.

For example, one can test if a linearly or exponentially decreasing speciation rate function is contained within the congruence class. In principle, there are no limitations to the choice of specific hypotheses and we provide several examples in our vignette (https://afmagee.github.io/ACDC). This option is useful when a researcher has a specific hypothesis regarding when diversification rates could have changed and what shape the diversification rates function might have.

In our primate HSMRF analysis, we inferred that the speciation rate changed abruptly in the last few million years, but the extinction rate remained comparably constant (Fig. 2). We note that the speciation rate appears to drive the changes in the “observed” net-diversification rate, i.e., in our originally inferred net-diversification rate. As an illustration, we explore here the alternative scenario if it instead was the extinction rate that drove the changes in the net-diversification rate. Because the net-diversification rate is defined as *δ*(*t*) = *λ*(*t*) − *μ*(*t*), we construct our new hypothesis for the extinction rate function as *μ*^′^(*t*) = *μ*_0_ − *λ*(*t*) where *μ*_0_ is any arbitrary value with *μ*_0_ ≥ *sup*(*λ*(*t*)) to ensure that *μ*^′^(*t*) ≥ 0. In `ACDC`, we only need to specify the alternative extinction rate functions *μ*^′^(*t*) and then call `congruent.models`.

~~~
mu_scaling <- c(1.1, 1.2, 1.5, 2.0)
mu_0s <- max(lambda(my_model$times)) * mu_scaling
mu_prime <- list()
for (i in seq_along(mu_scaling)){
mu_prime[[i]] <- local({
mu_0 <- mu_0s[i]
function(t) mu_0 - lambda(t)
})
}
alt_models <- congruent.models(my_model, mus = mu_prime)
plot(alt_models)
~~~

Even though we constructed the extinction rate functions to be responsible for the changes in the net-diversification rates, we observe that the corresponding speciation rate functions of the same congruence class are never constant (Fig. 3). This results in somewhat different net-diversification rate functions although both recent rate increases remain.

**Figure 3:**
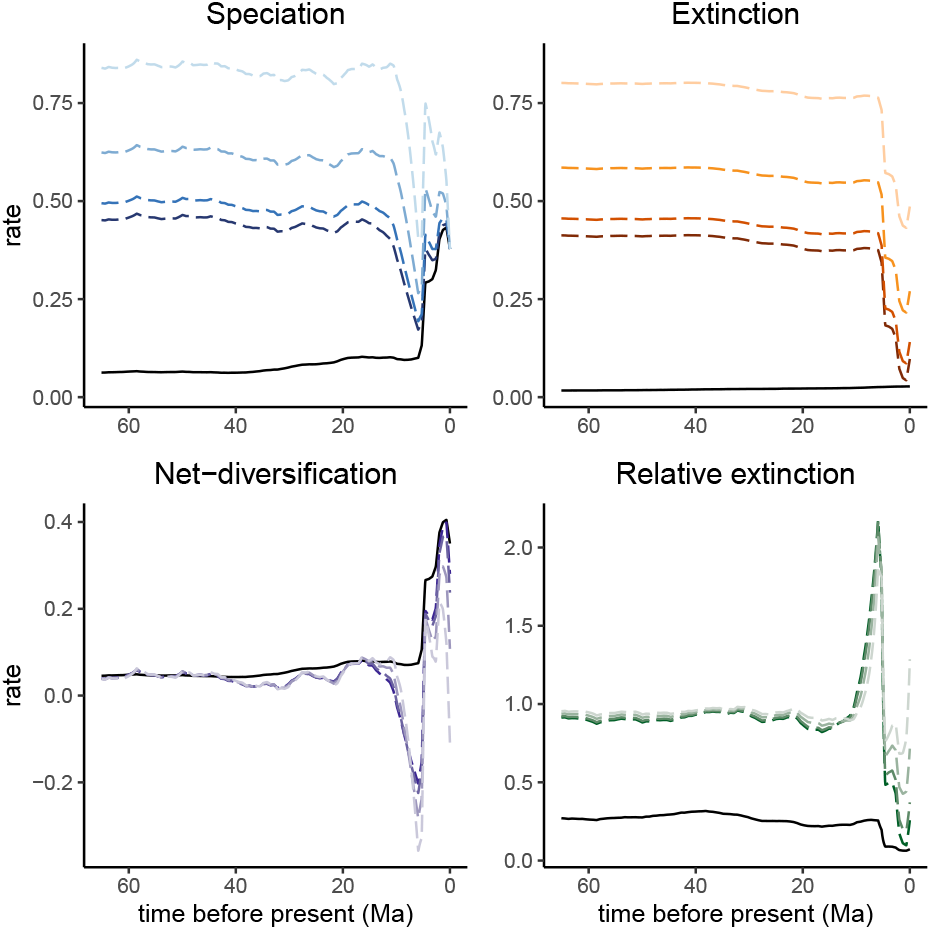
Alternative models contained in the congruence class. The solid black line depicts the *reference* model (*i.e.*, the model which we inferred and provided to `ACDC`). The four dashed lines in each plot show four different examples where we assumed that the extinction rate function was driving the net-diversification rates. The speciation rate functions, net-diversification rate functions and relative extinction rate functions are computed given the extinction rate functions to enforce that the models remain in the congruence class.

### 2.5 Exploring all congruent models, i.e., the congruence class

A second option is to explore the full congruence class. However, because there are infinitely many rate functions contained within the congruence class, we cannot explore every single model within the congruence class. Instead, we can sample randomly from a distribution of rate functions within the congruence class. We only need to specify how to create a random sample. For example, we could sample from different exponentially decreasing rate functions, or we could sample from a (discretized) Brownian motion.

In `ACDC`, we provide ways to randomly sample from several flexible rate distributions.

These distributions can be specified using the function `sample.basic.model`, and Table 1 provides an overview of the options for this function. A more detailed description is provided in our vignette (https://afmagee.github.io/ACDC).

**Table 1:**
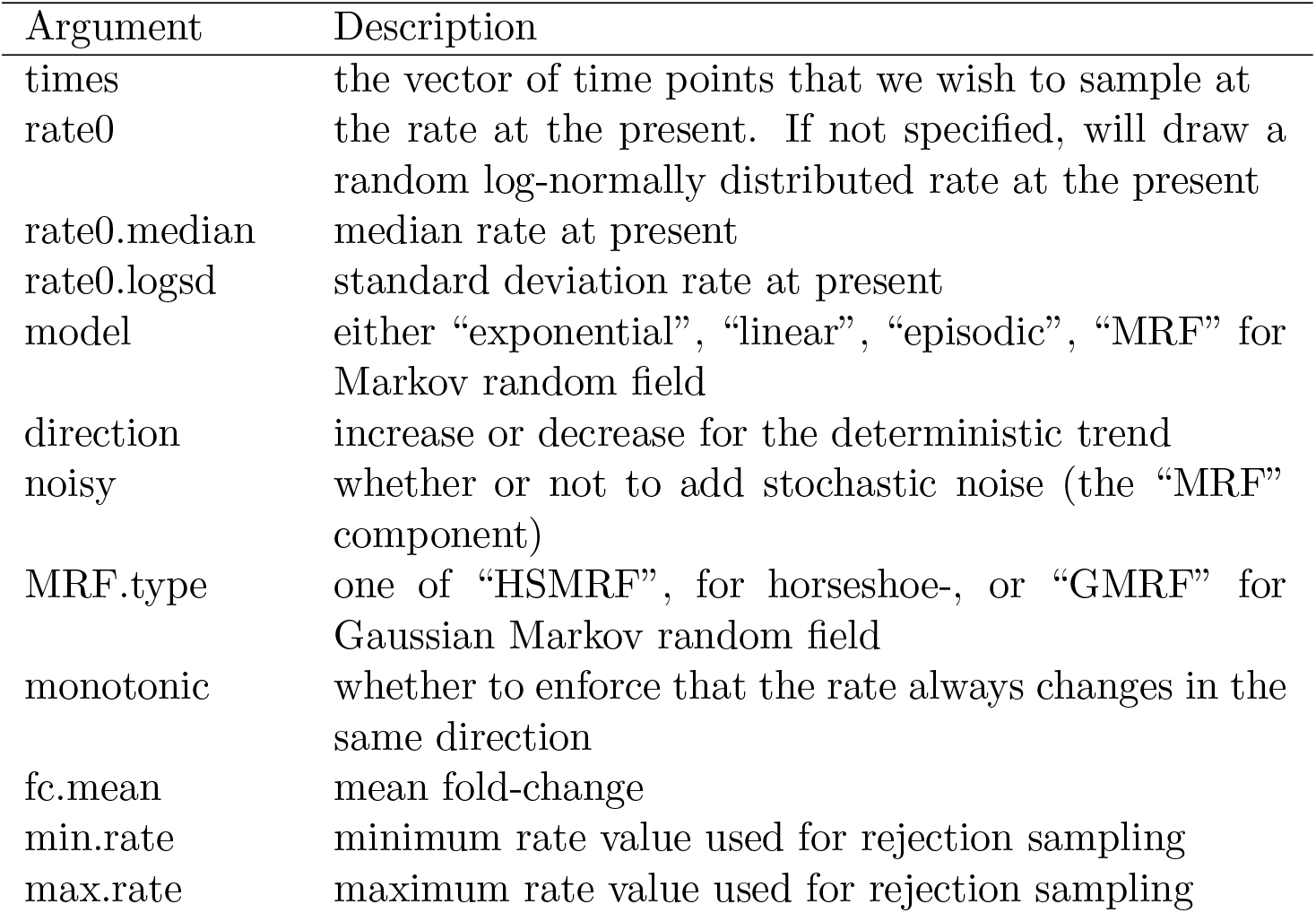
Options for sampling basic models using sample.basic.model.

As an example, we assume that an alternative extinction rate function corresponds to a Brownian motion. As a starting point for the Brownian motion at the present time *t*_0_ = 0, we sample 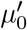 from a lognormal distribution. The distribution is centered around the reference estimate *μ*_0_, and we select the variance such that the central 95%-quantile of 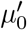 spans two orders of magnitude on the rate scale. Each preceding *μ_i_* is distributed as lognormal(*μ*_*i*−1_, *σ*), where *σ* ∼ HalfCauchy(0*, ζ*), and *ζ* is chosen depending on the number of epochs, so that the expected number of effective shifts in the rate is ln(2) (Magee *et al.* 2020). We repeat the entire procedure to draw each rate function.

~~~
extinction_rate_samples <- function() {
sample.basic.models(
times = primates_ebd[["time"]],
model = "MRF",
MRF.type = "GMRF",
max.rate = 1,
rate0.median = mu(0.0))
}
~~~

Then, we use this function—which specifies how to generate new samples for the extinction rate function— to sample from the congruence class, which is done using the function `sample.congruence.class`. Here, we sample 20 new extinction rate functions and automatically compute the corresponding speciation rate functions.

~~~
samples <- sample.congruence.class(
my_model,
num.samples=20,
rate.type="extinction",
sample.extinction.rates=extinction_rate_samples)
~~~

We can plot the HSMRF-samples in `ACDC`, using `plot(samples)` (Fig. 4, left and middle columns).

**Figure 4:**
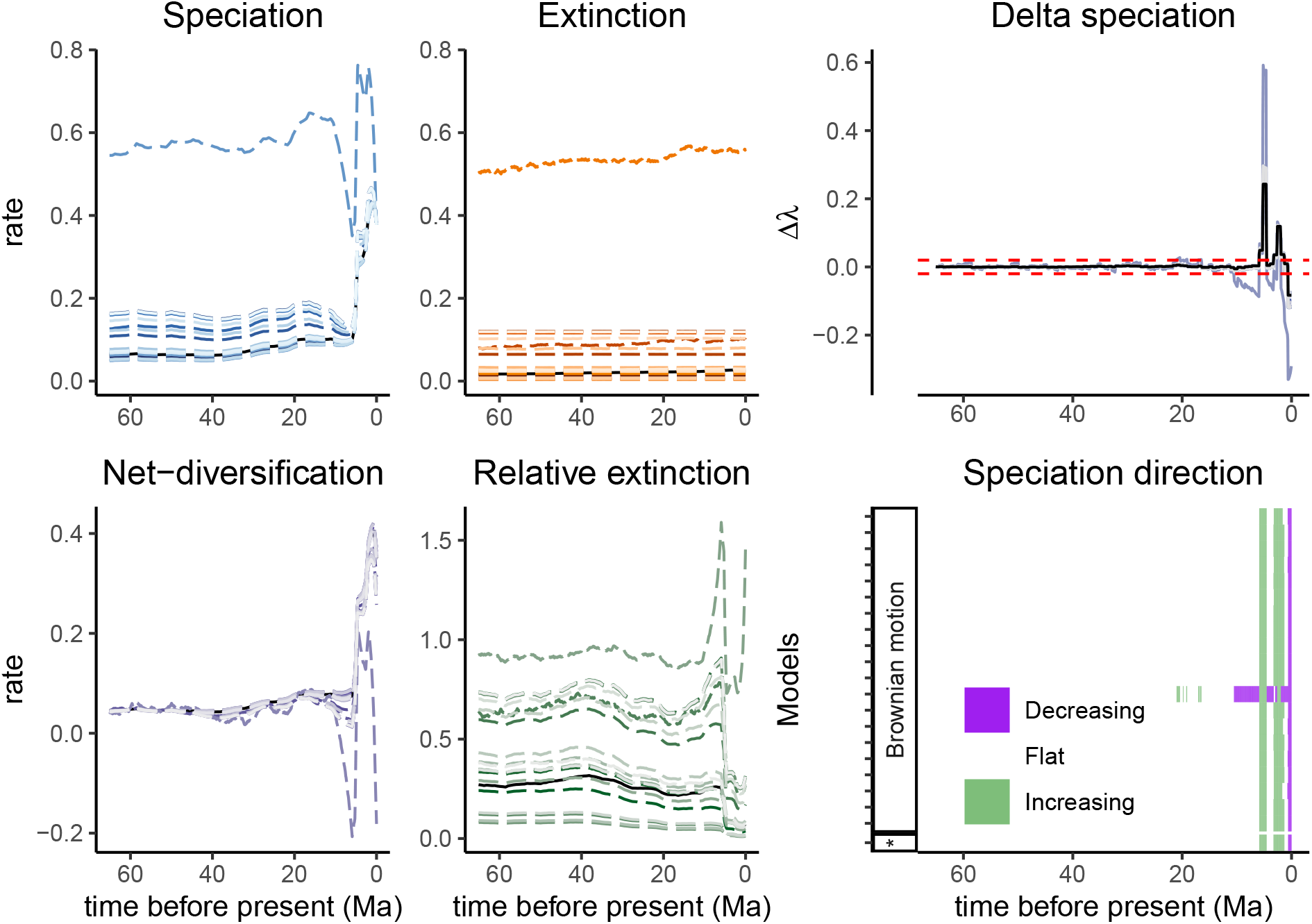
Congruent models where the extinction rates were sampled from a Brownian motion. The speciation rates were computed to match the extinction rates so that the models remain within the same congruence class. The net-diversification rates and relative extinction rate functions result from the speciation and extinction rate functions. The right column depicts the slope of the speciation rate (Δ*λ* = (*λ*_*i*−1_ − *λ_i_*)/Δ*t*), and a summary indicating whether the speciation rate function is decreasing, flat or increasing assuming a threshold for Δ*λ* of *±ϵ* = 0.02. The asterisk (*), and the solid black lines represent the *reference model.*

### 2.6 Summarizing trends over congruent models

Once several models from the congruence class are obtained, we compute summaries of our sampled rate functions. Recall that we are primarily interested in changes in diversification rates over time for these models. Therefore, we compute the amount of rate change within a small interval Δ*t* (*i.e.*, the slope of the rate functions)

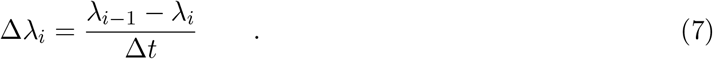

Next, we extract at which times the change in diversification rate (Δ*λ*) is larger than a pre-defined threshold *ϵ*: if Δ *λ* is larger than *ϵ* then we paint this time as a rate increase, and if Δ*λ* is smaller than −*ϵ* then we paint this time as a rate decrease. Alternatively, we can compute the normalized slope of the rate function:

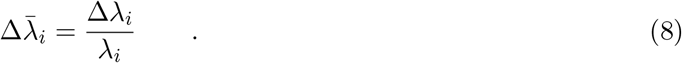

The absolute slope represents the change in the speciation rate per time, while the normalized slope represents the fold-change in the speciation rate per time. The absolute slope has the advantage that we can easily specify a comparable threshold over the entire diversification history, while the normalized slope has the advantage that we can easily specify a threshold that is comparable between analyses/datasets.

Outliers can signal single rate changes detected by Δ*λ* but could also be noise. Therefore, we implemented an option to smooth trends (either increases or decreases) by removing singleton outliers. We define an outlier as a time interval where both neighbors share the same trend but the interval itself has a different trend. The outlier is then replaced with the same trend as both its neighbors. This smoothing can clarify the overall signal to detect the total number of directional changes. However, smoothing might delete sharp or instantaneous rate changes.

In `ACDC`, we summarize the directional changes in the congruence class by specifying a threshold *ϵ*, in units of rate change per time.

~~~
summarize.trends(samples, threshold = 0.02)
~~~

The summary provides us with an overview of the trends: how many of the sampled models have speciation rate functions that were increasing or decreasing at a given time. For the example primate dataset, we observe that sampled models had two speciation rate increases very recently, and one additional speciation rate decrease at the present (Fig. 4). A more detailed description of how to summarize and interpret trends is provided in our vignette (https://afmagee.github.io/ACDC).

### 2.7 Accommodating uncertainty in rates

In the above example, we explored the congruence class for the posterior median diversification rates. We can also explore congruence classes for samples from posterior distributions. That is, as an example, we compute the congruence class for 20 samples from the posterior distribution and draw 20 alternative rate functions for each posterior sample. First, we load our posterior samples.

~~~
data(primates_ebd_log)
posterior <- read.RevBayes(
primates_ebd_log,
n_times = 1000,
max_t = 65,
n_samples = 20)
~~~

We plot the speciation rate functions from the posterior sample in the left panel of Fig. 5 (see Supplementary Scripts for information about plotting). Next, we generate congruent models for each of the samples from the posterior.

**Figure 5:**
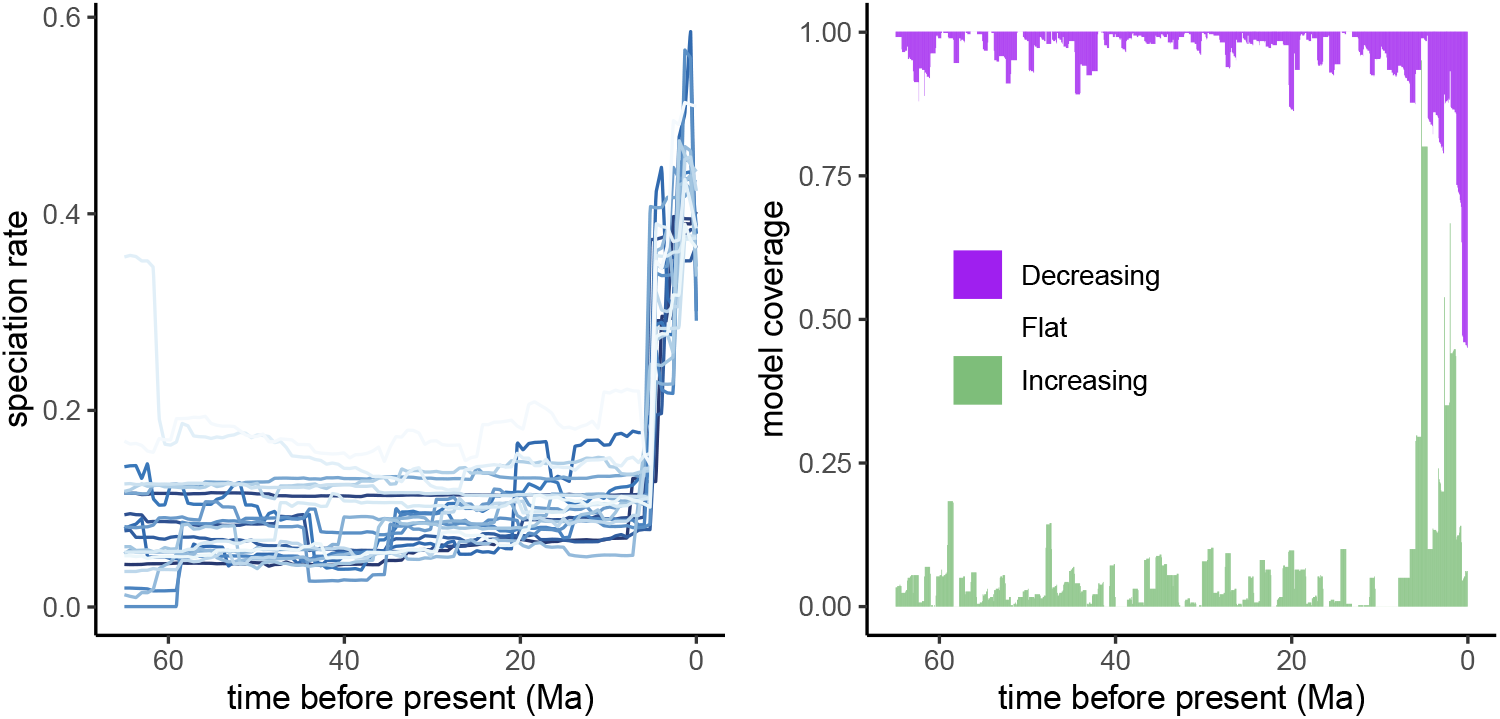
Left column: twenty posterior samples from the primates analyses. Right column: summary of trends over the posterior samples and congruence class samples for each posterior sample.

~~~
samples <- sample.congruence.class.posterior(
posterior,
num.samples = 20,
rate.type = "extinction",
model="MRF",
MRF.type = "GMRF",
max.rate = 3.0,
rate0.median = mu(0.0))
~~~

This selection yields 20 samples from the posterior, which are not congruent, but for each 20 samples we generated a subset of 20 additional congruent models. Next, we can summarize and plot the directions of change in the speciation rate function.

~~~
summarize.posterior(samples, threshold = 0.02)
~~~

Directional changes are summarized to show the number of models with an increase (or decrease, Fig. 5, right panel), rather than to keep results from the same model consistent across rows. In contrast to results based solely on the posterior median model (Fig. 4), the posterior samples show more disagreement between trends in direction of rate changes. Nevertheless, we observe general agreement among rate functions for the same three rate changes; two speciation rate increases in the last eight million years followed by one speciation rate decrease very close to the present.

### 2.8 Summary of available functions

Finally, we present an overview of the most important functions available in `ACDC` (Table 2). We demonstrated the core functionality in the previous sections, and we also provide a vignette where we explore more detailed features of the package. We designed `ACDC` to have few, but generic functions, that allow for flexibility in exploration of the rate space in the congruence class.

**Table 2:** A summary of the core functions used in ACDC.

| Function | Description |
| --- | --- |
| `congruent.models` | constructs models that are congruent with a reference model |
| `create.model` | creates an **ACDC** model object |
| `model2df` | converts an **ACDC** model object to a data frame, e.g. for plotting |
| `plot.ACDC` | plots a birth-death model |
| `plot.ACDCset` | plots a set of congruent models |
| `sample.basic.models` | samples rate functions from various distributions (see Table 1) |
| `sample.congruence.class` | samples models from the congruence class |
| `sample.congruence.class.posterior` | sample congruence class from the posterior |
| `sample.rates` | samples custom rate functions |
| `summarize.posterior` | plots a summary of the directional trends in the posterior |
| `summarize.trends` | plots a summary of the directional trends in the congruence class |
| `read.RevBayes` | reads a RevBayes log file |

We designed `ACDC` to use standard ggplot objects (Wickham 2016) so that plots can easily be manipulated as any other ggplot objects. For example, it is possible to change the axis limits, axis labels, or the time scale. This allows for flexibility in visualizing the congruence class for other data sets than we have exemplified here.

## 3 Discussion and conclusions

In this paper we present the R package `ACDC`, Analysis of Congruent Diversification Classes. `ACDC` is available on CRAN and the source code is available from GitHub (https://github.com/afmagee/ACDC). `ACDC` enables easy testing of the impact of non-identifiable diversification rates. Specifically, with `ACDC` anyone can test specific alternative diversification rate hypotheses (Morlon *et al.* 2020), or explore equally probable diversification rate models for shared characteristics. Thus, non-identifiability of diversification rates can be incorporated into conclusions about the process shaping historical biodiversity.

In our main example of exploring a congruence class, we sampled alternative rate functions from a Brownian motion process. This choice reflects our belief that Brownian motion might be a good approximation of how diversification rates have changed over time, but other approaches should be considered (see Condamine *et al.* 2018, Magee *et al.* 2020 and Palazzesi *et al.* 2022 for comparisons of diversification rate models through time). To assist with this, `ACDC` provides functions to generate alternative rate functions with stochastic changes, as well as diversification rate functions with deterministic trends (e.g., exponential and linear). The existing functions to explore the congruence class can accept any type of rate functions. It remains unclear what the best approach to sample alternative rate functions is, and we leave the decision to the researcher for the specific study.

The primary output of `ACDC` is the congruence class and models belonging to this congruence class. Since changes in diversification rates are generally of interest, rather than the rates themselves, we focus on summaries of the congruence class showing directional trends in diversification rates (increasing, decreasing or flat). However, these summaries strongly depend on the chosen threshold for assessing whether the diversification rate was changing or not. We chose an arbitrary threshold of *ϵ* = 0.02, which signifies that any rate change of ±0.02 per million years is a significant trend. We recommend in practice testing summaries using a variety of thresholds. We strongly recommend researchers begin by exploring and visualizing specific models (as in Section 2.4), before examining broader sections of the congruence class (as in Sections 2.5 and 2.6).

Finally, non-identifiability of diversification rates extends to diversification rates inferred from phylogenies with fossil taxa (Louca *et al.* 2021). In `ACDC`, we only focus on speciation and extinction rate functions and omit fossilization rates. However, it is possible to analyze a congruence class obtained from a phylogeny with fossil taxa using an approach analogous to what we described here. If a specific new extinction rate function was chosen, then both the speciation rate function and fossilization rate function can be computed given the congruence class. The results with `ACDC` are still valid for speciation and extinction rates, and could be considered as if the fossilization was simply not shown.

Non-identifiability of diversification rates has questioned the reliability and interpretation of diversification rate estimates from current approaches. Non-identifiability is a very real problem and should not be neglected. `ACDC` provides a new tool to assess if the conclusions drawn about patterns of diversification rates are robust despite the existence of infinitely many alternative diversification models.

## 4 Acknowledgments

We thank Luiza Fabreti, Ronja Billenstein and Killian Smith for feedback about visualizing and interpreting congruence classes.

## 5 Funding

This work was supported by the Deutsche Forschungsgemeinschaft (DFG) Emmy Noether-Program (Award HO 6201/1-1 to S.H.). AFM was partially supported by National Science Foundation grant DGE-1762114 and National Institutes of Health grant R01 AI153044.

## 6 Author Contributions

SH and AFM conceived the study. SH, BTK and AFM developed the R package `ACDC`. SH and BTK drafted the manuscript. All authors revised and approved the manuscript.

## Notes

https://afmagee.github.io/ACDC/overview.html

### Competing Interest Statement

The authors have declared no competing interest.

