## Supplementary Material for "ACDC: Analysis of Congruent Diversification Classes"

#### Supporting Information

Sebastian Höhna[1,2*], Bjørn T. Kopperud[1,2] and Andrew F. Magee[3]

[1] *GeoBio-Center, Ludwig-Maximilians-Universität München, Richard-Wagner Straße 10, 80333 Munich, Germany*
[2] *Department of Earth and Environmental Sciences, Paleontology & Geobiology, Ludwig-Maximilians-Universität München, Richard-Wagner Straße 10, 80333 Munich, Germany*
[3] *Department of Human Genetics, University of California, Los Angeles, 90095, U.S.A.*

### Contents

S1 Derivation of speciation rate function within the congruence class . . . . . . . . . . . . . . . . 2
S2 Accuracy of the piecewise linear method . . . . . . . . . . . . . . . . . . . . . . . . . . . . . . . . 3

### S1 Derivation of speciation rate function within the congruence class

The pulled net-diversification (Louca & Pennell, 2020) is one of the identifiable parameters in the reconstructed birth-death model. It is defined as a differential equation

$$r_p(t) = \lambda(t) - \mu(t) + \frac{1}{\lambda(t)}\frac{d\lambda(t)}{dt} \quad . \tag{S1}$$

The pulled rates are calculated automatically in `ACDC` for any model object. Recall that the pulled net-diversification rate, together with $\lambda(0)$, completely characterizes the congruence class. In order to construct a congruent model, we can propose an alternative rate function $\mu'(t)$, and solve for $\lambda(t)$. In the main text, we used a difference equation to approximate the pulled net-diversification rate as

$$r_p(t_i) = \lambda_i - \mu_i + \frac{1}{\lambda_i}\frac{\lambda_i - \lambda_{i-1}}{\Delta t} \quad . \tag{S2}$$

The initial $\lambda_0$ at the present is equal for all models in the congruence class. Next, we need to calculate $\lambda_i$ for each preceding time $i$ in the rate function. We would like to solve for $\lambda_i$ using Eq. (S2). In the following steps we will detail the steps needed to solve for $\lambda_i$. First, we make a common denominator $\lambda_i \Delta t$

$$r_p(t_i) = \frac{\lambda_i^2 \Delta t - \mu_i \lambda_i \Delta t + \lambda_i - \lambda_{i-1}}{\lambda_i \Delta t} \quad . \tag{S3}$$

We multiply both sides by $\lambda_i \Delta t$

$$r_p(t_i)\lambda_i \Delta t = \lambda_i^2 \Delta t - \mu_i \lambda_i \Delta t + \lambda_i - \lambda_{i-1} \quad . \tag{S4}$$

We collect by $\lambda_i$, and rearrange

$$\lambda_i^2 \Delta t + \lambda_i(1 - \mu_i \Delta t - r_p(t_i)\Delta t) = \lambda_{i-1} \quad . \tag{S5}$$

We divide by $\Delta t$

$$\lambda_i^2 + \lambda_i \frac{(1 - \mu_i \Delta t - r_p(t_i)\Delta t)}{\Delta t} = \frac{\lambda_{i-1}}{\Delta t} \quad . \tag{S6}$$

Since we have both $\lambda_i^2$ and $\lambda_i$, we want to use quadratic factorization: $(\lambda_i + a)^2 = \lambda_i^2 + \lambda_i 2a + a^2$, with $a = (1 - \mu_i \Delta t - r_p(t_i)\Delta t)/(2\Delta t)$. To complete the square, we add $a^2$ to both sides

$$\lambda_i^2 + \lambda_i 2a + a^2 = \frac{\lambda_{i-1}}{\Delta t} + a^2 \quad , \tag{S7}$$

$$(\lambda_i + a)^2 = \frac{\lambda_{i-1}}{\Delta t} + a^2 \quad . \tag{S8}$$

Next, we substitute back $a$

$$\Big(\lambda_i + \frac{1 - \mu_i \Delta t - r_p(t_i)\Delta t}{2\Delta t}\Big)^2 = \frac{\lambda_{i-1}}{\Delta t} + \Big(\frac{1 - \mu_i \Delta t - r_p(t_i)\Delta t}{2\Delta t}\Big)^2 \quad . \tag{S9}$$

Then, we take the square root on both sides, and leave $\lambda_i$ on the left side

$$\lambda_i = \sqrt{\frac{\lambda_{i-1}}{\Delta t} + \frac{(1 - \mu_i \Delta t - r_p(t_i)\Delta t)^2}{4(\Delta t)^2}} - \frac{(1 - \mu_i \Delta t - r_p(t_i)\Delta t)}{2\Delta t} \quad . \tag{S10}$$

Finally, we arrange yo obtain our main solution,

$$\lambda_i = \frac{\sqrt{4\lambda_{i-1}\Delta t + (1 - \mu_i \Delta t - r_p(t_i)\Delta t)^2} + \mu_i \Delta t + r_p(t_i)\Delta t - 1}{2\Delta t} \quad . \tag{S11}$$

#### S2 Accuracy of the piecewise linear method

We use a piecewise linear method to approximate the rate functions. The idea is that, for sufficiently many pieces, or sufficiently small $\Delta t$, we can approximate any continuously varying rate function. How many pieces is considered sufficient? To assess this question, we require some sort of benchmark. One property of the congruence class is that all models must have the same deterministic lineage-through-time (dLTT) curve. Thus, we can check if our transformation to another model within the congruence class worked. We used the R-package TESS (Höhna *et al.*, 2016) to calculate the dLTT for a selection of models with different number of linear pieces (Fig. S1). In the original analysis, we used 100 epochs/intervals to estimate the piecewise constant diversification rates. It appears that 10, 20, and even 50 pieces are inadequate. In our view, 500 or more pieces are sufficient for precisely setting up the primates congruence class, and for proposing alternative congruent models.

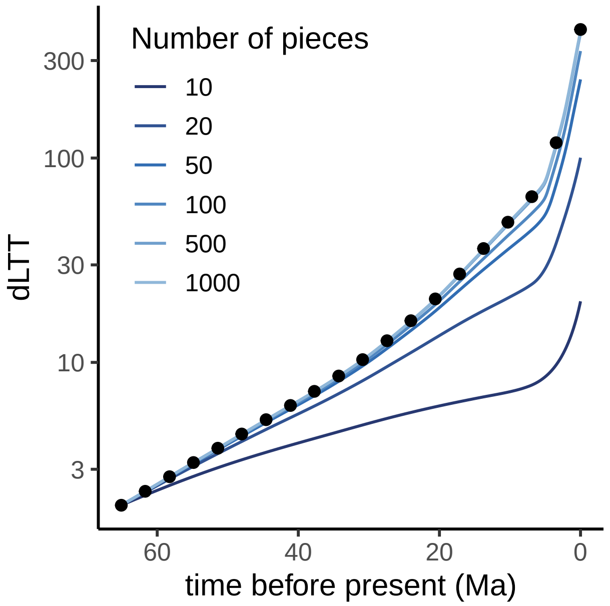


**Figure S1:** The deterministic lineage-through-time (dLTT) for two congruent models generated using a range of number of linear pieces. If the transformation worked, i.e., we obtained truly another diversification rate model within the same congruence class, then the dLTT must be identical. We computed the dLTT curves using TESS (Höhna *et al.*, 2016). The dotted line represents the reference model (i.e. the dLTT based on the posterior median speciation and extinction rate estimates for the Primates phylogeny), where we used 1000 linear pieces. The solid lines depict congruent models where we proposed a new extinction rate $\mu'(t) = 2 \times \sup(\lambda(t)) - \lambda(t)$, and inferred the congruent speciation rate. The proposed models approach the dLTT of the reference model as the number of pieces increase. 500 and 1000 pieces are nearly indistinguishable.
